# Genetic diversity and population structure of razor clam *Sinonovacula constricta* in Ariake Bay, Japan, revealed using RAD-Seq SNP markers

**DOI:** 10.1101/2020.10.27.358150

**Authors:** Ryo Orita, Yukio Nagano, Yoshio Kawamura, Kei Kimura, Genta Kobayashi

## Abstract

The razor clam *Sinonovacula constricta* is a commercially important bivalve in Japan. The current distribution of this species in Japan is limited to Ariake Bay, where the fishery stock is declining. It is necessary to understand the genetic population structure of this species in order to restore the fishery stock while preserving the genetic diversity of the clam. Here, we report for the first time the genetic population structure of *S. constricta* in Ariake Bay, Japan. Paired-end restriction site-associated DNA sequencing (RAD-Seq) analyzed samples of *S. constricta* collected from seven mudflats located along Ariake Bay. Two different genetic populations exist in Ariake Bay, one inhabiting wild habitats and the other inhabiting the transplanted area of artificial seedlings. Our results suggest that genetic differentiation occurred between these two populations (*F*_*st*_ value = 0.052), and a high level of genetic differentiation is maintained between the two groups. In the future, monitoring the interbreeding status of the two genetically distinct populations and the genetic differentiation within each population is important for conserving the genetic diversity of *S. constricta* in Japan.

## Introduction

The technique of artificially producing seedlings and releasing them into fishing grounds is an effective method of aquaculture enhancement. This technique has been implemented with a variety of fishery species (e.g., finfish [1, 2], flounder [3], shellfish [4-7]). In artificial seed production, large numbers of seedlings are often produced by a few broodstocks and released into natural seawater, which reduces the genetic diversity of wild populations in the field. Therefore, when seedlings are produced and released into natural seawater, there is a need to understand the genetic structure within the wild population in order to preserve genetic diversity.

The razor clam *Sinonovacula constricta* (Lamarck, 1818) is an important economic species in East Asian coastal waters [8-10] and is widely farmed in China [10]. This species was once distributed from central to western Japan [11-12] but is now restricted to the mudflats around Ariake Bay on the west coast of Kyushu in western Japan. In Japan, razor clams are not farmed, and the catch depends on the amount of resources in natural seawaters. According to a report by Ministry of the Environment, Japan [13], the annual catch of *S. constricta* in Ariake Bay often recorded more than 1000 tons per year from the 1900s to the early the 1920s, but it decreased to less than 500 tons in most years from the latter half of the 1920s to the latter half of the 1980s, and no catches have been made since 1994 because of the marked decline in the abundance of this species. In order to restore the *S. constricta* resources, the Saga Prefectural Ariake Fisheries Research and Development Center has developed seed production techniques [14, 15], and since 1988, artificial seedlings of *S. constricta* have been released into the transplanting areas established in natural seawaters [16] and have been conserved in transplanting areas as populations to produce subsequent generations. However, the genetic population structure of *S. constricta* within Ariake Bay has not been examined.

In recent years, technological innovations have made the genome-wide analysis of non-model organisms more widely available, and these techniques are useful for examining the genetic population structure of non-model aquatic organisms [17-20]. In the present study, to clarify the genetic population structure of *S. constricta* in Ariake Bay, we conducted field sampling surveys at 11 mudflats located along the bay in 2019 (Fig. 1) and used restriction-site-associated DNA sequencing (RAD-Seq) to analyze population genetics and phylogenetics. We propose guidelines concerning the optimal seed production and seed release methods for the conservation of the genetic diversity of *S. constricta* in Japan.

**Figure 1.**
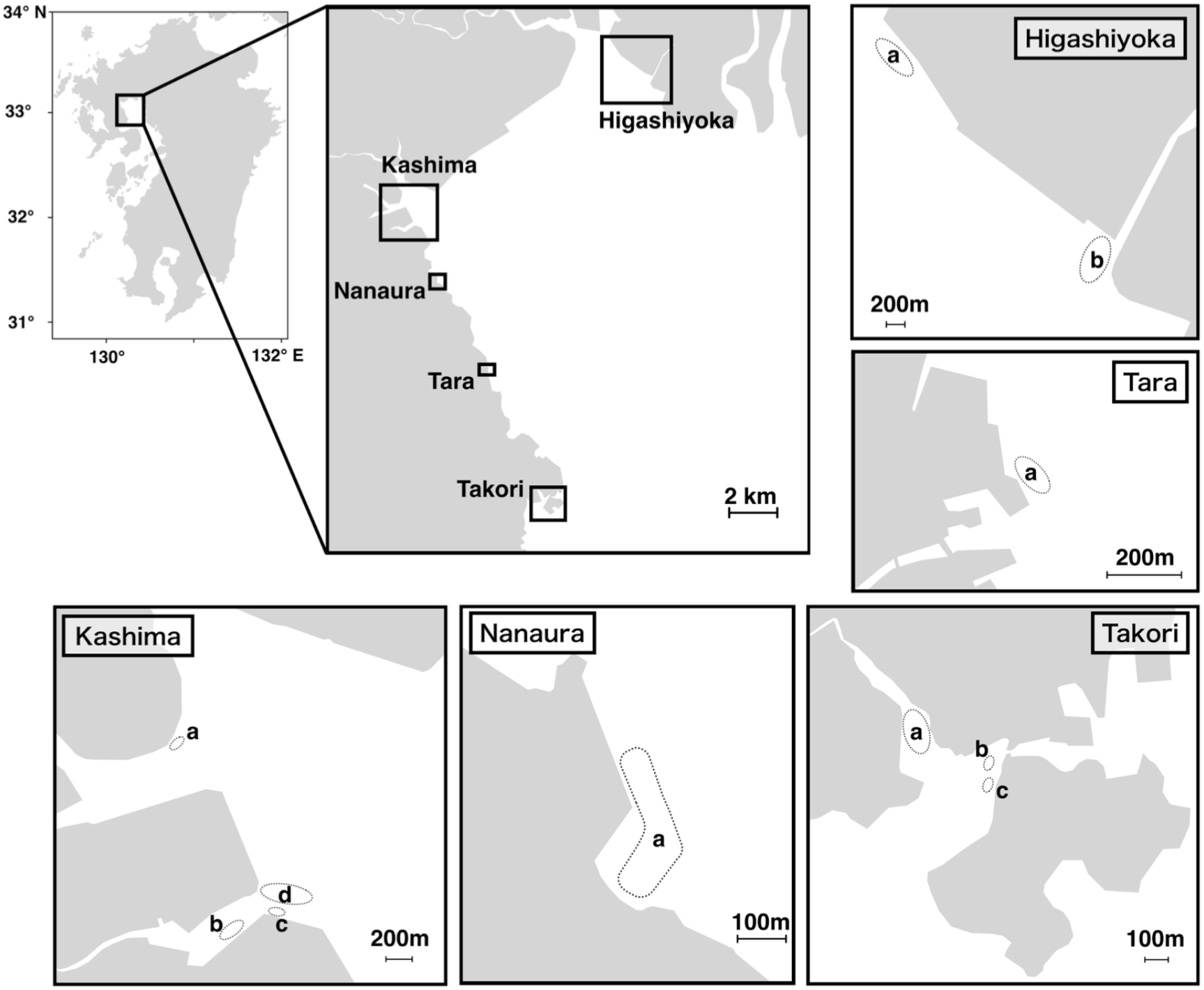
Sampling sites. The dotted line shows the area explored.

## Results

### Sampling data

We collected 28 razor clams from seven of the 11 mudflats and found no razor clams on the remaining four mudflats (Table 1, Fig. 1). The mean CPUE of the seven mudflats where individuals could be collected was 3.2 (± 3.0 SD) ind. person^−1^ hour^−1^, and the range of CPUE at each site was 0.3 to 8.5 ind. person^−1^ hour^−1^. The shell length of the collected individuals was 39.7 to 94.6 mm, implying that they contained specimens from 1 to 5 years old. The mean shell length of individuals collected from the transplanted area was 86.0 (± 4.5 SD) mm for the 3-year-old seedlings produced in 2016 (i.e., sample ID: 2016.1-2016.7) and 92.9 (± 1.1 SD) mm for the 5-year-old seedlings produced in 2014 (i.e., sample ID: 2014.1-2014.5).

**Table 1.** Sampling information for *Sinonovacula constricta* in the Ariake Bay, Japan

| Sampling site | Sampling day | Habitat type | CPUE*<br>(ind. person<sup>-1</sup> h<sup>-1</sup>) | Sample ID | Shell length<br>(mm) |
| --- | --- | --- | --- | --- | --- |
|  | Nov. 11, 2019 | Wild | 0.3 | Kashima a.1 | 39.7 |
| Kashima_b | Oct. 11, 2019 | Wild | 1.3 | Kashima b.1 | 46.6 |
|  |  |  |  | Kashima b.2 | 45.2 |
|  |  |  |  | Kashima b.3 | 54.7 |
|  |  |  |  | Kashima b.4 | 54.0 |
| Kashima_c | Nov. 19, 2019 | Wild | 4.0 | Kashima c.1 | 44.7 |
|  |  |  |  | Kashima c.2 | 51.4 |
|  |  |  |  | Kashima c.3 | 48.1 |
|  |  |  |  | Kashima c.4 | 67.0 |
|  | Oct. 23, 2019 | Wild | N/A<sup>†</sup> | Kashima c.5 | 70.2 |
| Kashima_d | Nov. 11, 2019 | Wild | 0.0 | — | — |
| Nanaura_a | Nov. 13, 2019 | Wild | 0.7 | Nanaura a.1 | 47.0 |
|  |  |  |  | Nanaura a.2 | 53.9 |
| Tara_a | Nov.13, 2019 | Wild | 0.0 | — | — |
| Takori_a | Nov. 15, 2019 | Wild | 1.8 | Takori a.1 | 75.0 |
|  |  |  |  | Takori a.2 | 55.7 |
|  |  |  |  | Takori a.3 | 81.7 |
|  |  |  |  | Takori a.4 | 94.6 |
| Takori_b | Nov. 15, 2019 | Transplanted area<sup>‡</sup> | 5.5 | 2014.1 | 92.8 |
|  |  |  |  | 2014.2 | 94.8 |
|  |  |  |  | 2014.3 | 92.1 |
|  |  |  |  | 2014.4 | 91.9 |
|  |  |  |  | 2014.5 | 92.9 |
| Takori_c | Nov. 15, 2019 | Transplanted area<sup>‡</sup> | 8.5 | 2016.1 | 88.4 |
|  |  |  |  | 2016.2 | 87.0 |
|  |  |  |  | 2016.3 | 83.7 |
|  |  |  |  | 2016.4 | 82.6 |
|  |  |  |  | 2016.5 | 86.7 |
|  |  |  |  | 2016.6 | 79.7 |
|  |  |  |  | 2016.7 | 93.6 |
| Higashiyoka_a | Nov. 12, 2019 | Wild | 0.0 | — | — |
| Higashiyoka_b | Oct. 18, 2019 | Wild | 0.0 | — | — |
| N/A<sup>†</sup> | Dec. 10, 2006 | Transplanted area<sup>‡</sup> | N/A<sup>†</sup> | 2005.1 | 66.7 |
|  |  |  |  | 2005.2 | 74.7 |

\*: Catch per unit effort (individual person<sup>-1</sup> hour<sup>-1</sup>)
<sup>†</sup>: Not available
<sup>‡</sup>: Transplanted area of artificial seedlings

### RAD-Seq data

We used 30 individuals for RAD-Seq analysis, including 16 individuals collected from wild habitats, 12 individuals collected from the transplanted area of artificial seedlings, and 2 individuals stored at the Saga Prefectural Ariake Fisheries Research and Development Center (Table 1). In all, 401,109,332 raw reads were generated from the HiSeq X run, and 249,040,001 reads were retained after discarding reads with low quality and adapters. The final dataset included 137,246 loci, with an average of 679.38 bp (SE ± 0.18) of genotyped sites per locus.

### Population genetics structure

The genetic relationships among sampling sites were clearly divided into individuals collected from wild habitat and those from the transplanted area of artificial seedlings along the axis of PC1 (Fig. 2a). The PC2 axis represents the differences within each habitat type. In the PCA plot, the coordinates of the transplanted samples were more widely scattered than those of the wild samples were, showing more diverse genetic relationships among transplanted samples. When evaluated by shell length (Fig. 2b), genetic diversity was higher in the 75–90 mm size group than that in the younger age group with smaller shells. The results of admixture analysis also showed a significant genetic difference between the individuals collected from the wild versus those collected from the transplanted areas of artificial seedlings (Fig. 3). The *F*_*st*_ value between populations consisting of individuals collected from the wild and the transplanted areas was 0.052. Owing to the calculation of the pairwise *F*_*st*_values for each individual within the two populations, there was no significant difference in the mean values between these populations (Fig. 4). The pairwise *F*_*st*_ values within the population were more varied in the transplanted area (from 0.44 to 0.64) than those in the wild habitat (from 0.53 to 0.56). There were no significant differences in shell morphology between these two genetically different populations (Supplementary Fig. S1).

**Figure 2.**
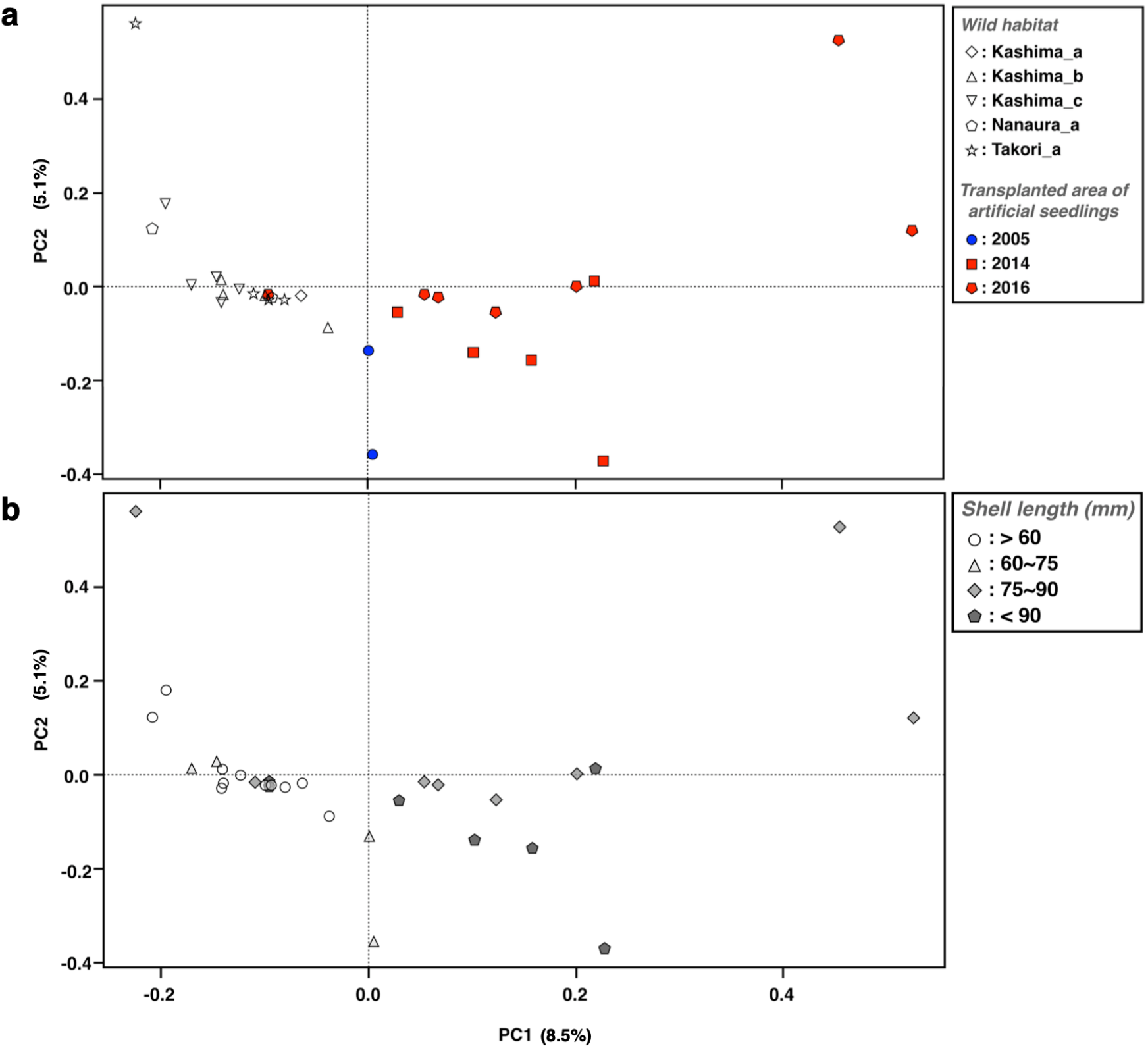
Biplots of principal component analysis. (a) Showing by sampling areas. (b) Showing by shell sizes.

**Figure 3.**
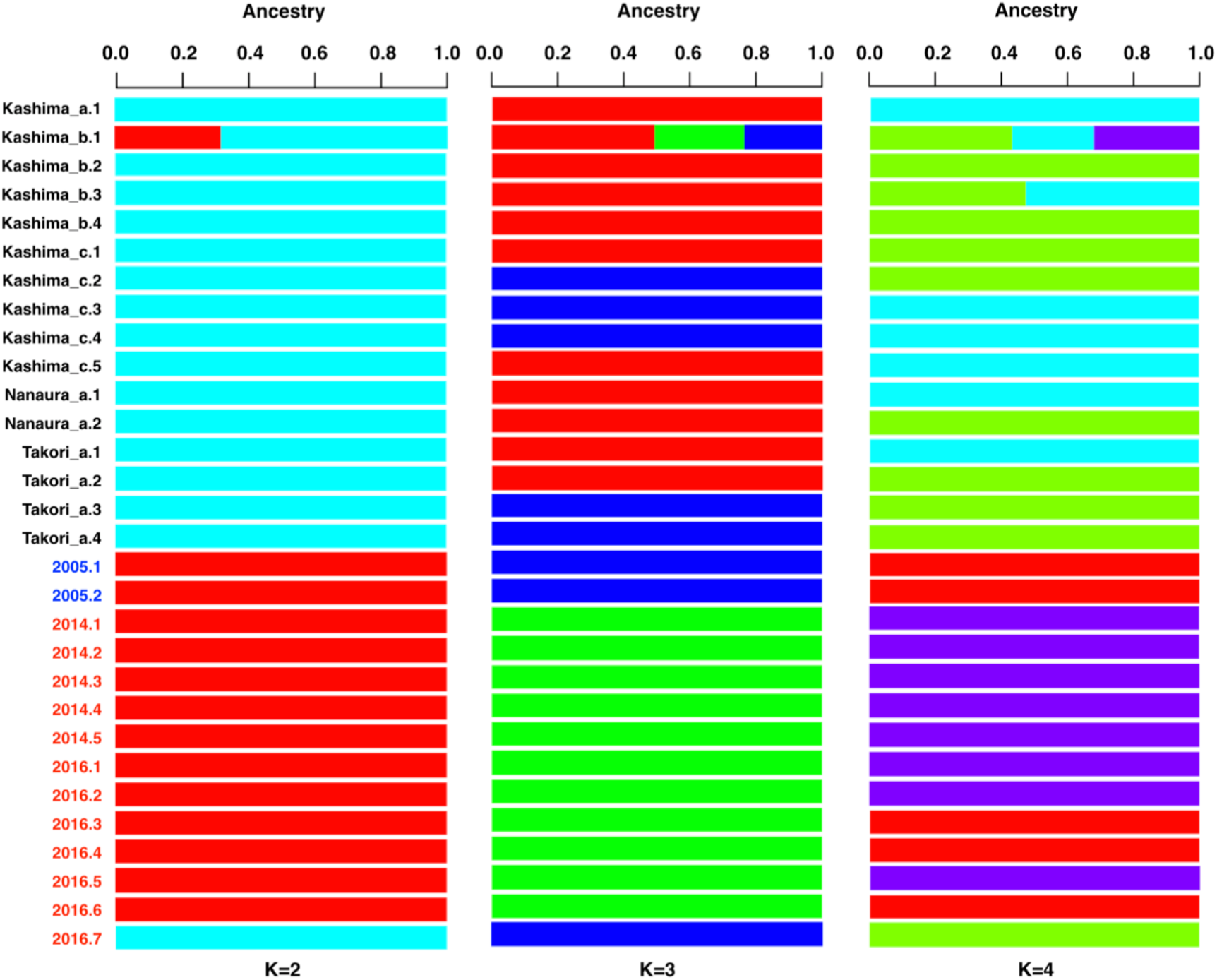
Admixture analysis of *S. constricta* in the Ariake Bay, Japan.

**Figure 4.**
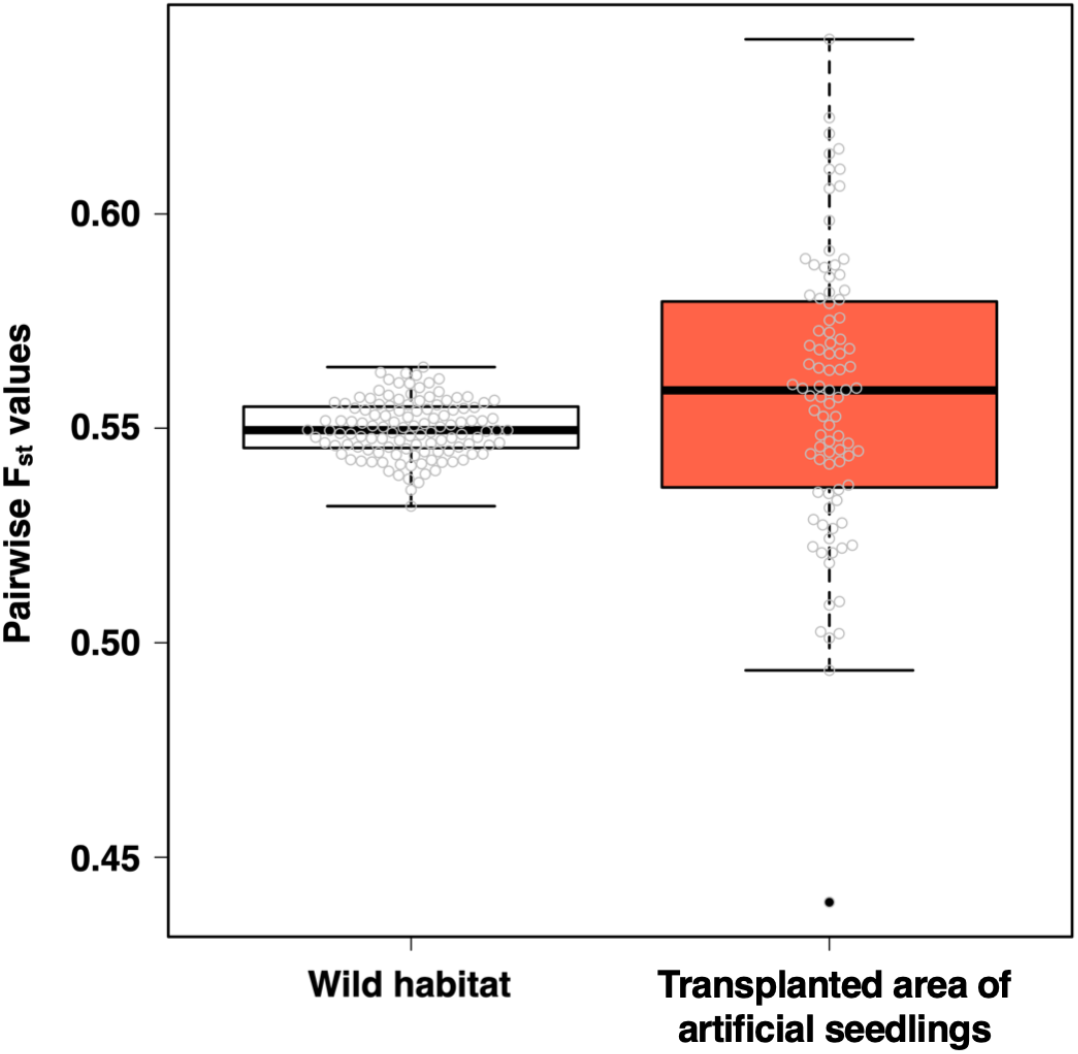
Pairwise F_st_ values between individuals in each population.

### Phylogenetic tree analysis

ML tree and SVDquartets analysis of RAD-Seq data showed that the phylogenies were highly consistent with each other (Fig. 5a and 5b). The obtained phylogenetic trees were clearly divided into individuals collected from the wild habitat and those collected from the transplanted area of artificial seedlings, with the exception of one sample (i.e., sample ID: 2016.7).

**Figure 5.**
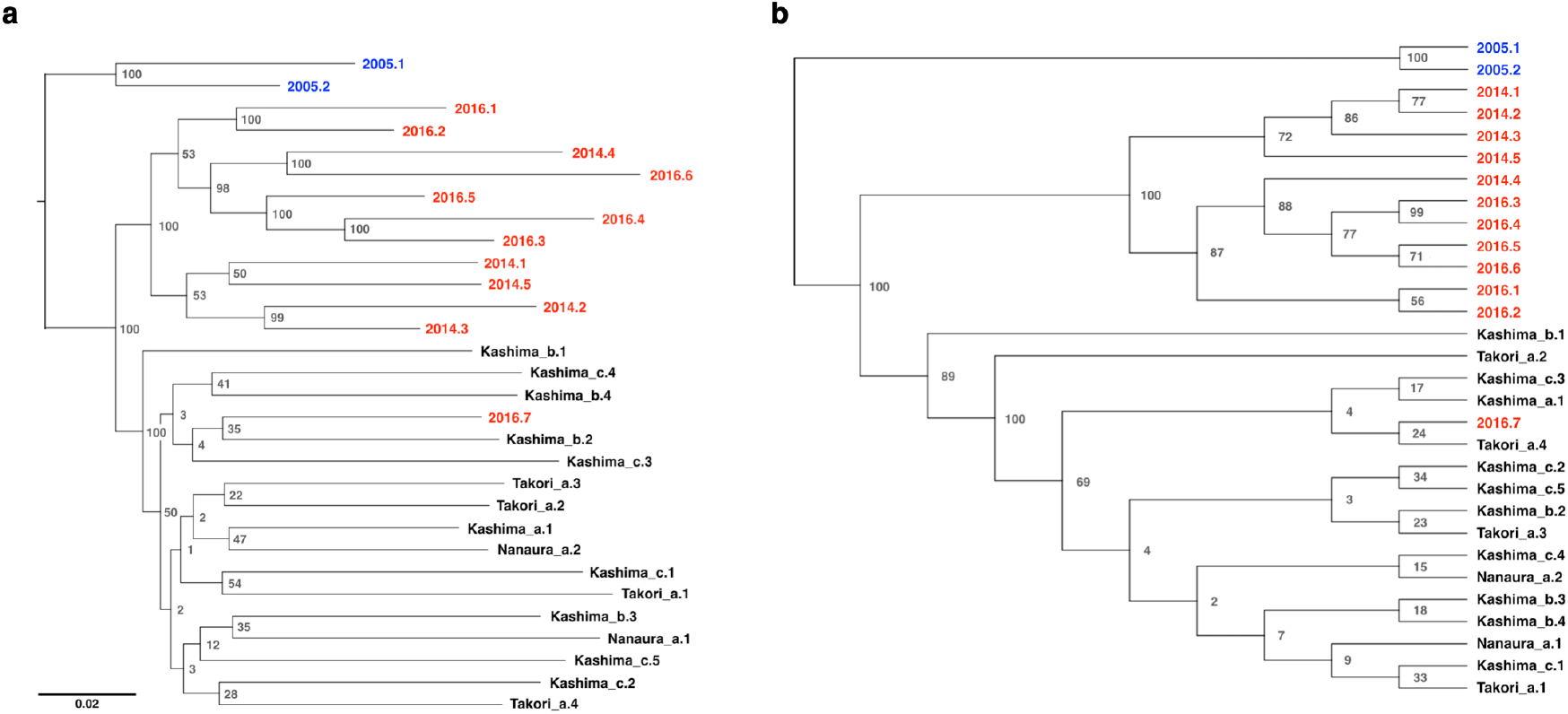
Phylogenetic trees of *S. constricta* inferred from RAD-Seq dataset (a) Maximum likelihood phylogeny estimated using RAxML. (b) Species tree inferred using SVDquartets. Numbers at the nodes indicate bootstrap values (1000 replicates).

## Discussion

### Genetic population structure of *S. constricta* in the Ariake Bay, Japan

All our analyses of the RAD-Seq data showed that individuals collected from wild habitats were genetically different from those collected from sites where artificial seedlings were transplanted. Only one sample collected from the area transplanted in 2016 (sample ID: 2016.7) showed different results from other seedlings collected in the same area, and belonged to a population consisting of individuals collected from the wild habitat (Fig. 3, Fig. 5a and 5b). Since this individual had a distinctly larger shell length than other 3-year-old seedlings in the same area, and its size was the same as that of 5-year-old seedlings (Table 1), it was likely a wild individual inhabiting the transplanted area prior to the 2016 transplantation. Therefore, we consider that there were two genetically distinct populations in Ariake Bay in 2019.

The reason for the differences between these two populations is the different parental origins of the seedlings. The parental origins can be divided into the Nanaura mudflat and the Kashima mudflat for a number of reasons. Artificial seedlings of *S. constricta* have been released by the Saga Prefectural Ariake Fisheries Research and Development Center into transplanting areas established in several mudflats along the coast of Ariake Bay. The seedlings collected from the transplanted area in this study (Takori_b and Takori_c) originated from parents at the Nanaura mudflat (hereinafter referred to as Nanaura lineage population), since the parents were collected from the Nanaura mudflat before 2010 and released into the transplanting areas of the Takori mudflat. In contrast, individuals collected from the wild habitat of Kashima, Nanaura, and Takori mudflats are likely to have originated from parents of Kashima mudflat (hereinafter referred to as the Kashima lineage population). In 2017 and 2018, the seedlings produced by parents collected from Kashima mudflat were released into new transplanting areas at Takori and Nanaura mudflats due to declining populations in wild and transplanted areas in these mudflats. Therefore, the 1-to 2-year old individuals collected in wild habitats in 2019 were likely the offspring of the transplanted Kashima seedlings in 2017 and 2018.

Artificial transplantation of aquatic organisms has been reported to alter the geographic genetic structure of wild populations [21-24], including that of *S. constricta* (in this case, along the Chinese coast) [25]. Anthropogenic transplantation likely strongly influenced the distribution of the genetic population structure of *S. constricta* in Japan. Comparison of the genetic distribution of species with a life history similar to that of *S. constricta* but those that are not exploited for fisheries can enhance our understanding of anthropogenic impacts on the genetic distribution of *S. constricta* in the Ariake Bay.

### Genetic differentiation among and within populations

For artificial seed production and release of seedlings, it is necessary to understand the distribution of populations with different genetic structures in natural seawater. Therefore, it is necessary to select broodstocks and release areas according to the degree of genetic differentiation among and within populations. *F*_*st*_ values are widely used as a measure of the degree of genetic differentiation among and within populations [26-28]. According to Niu et al. [25], the *F*_*st*_ value among populations of *S. constricta* at 10 sites along the coast of mainland China is 0.044. The *F*_*st*_ value between the two populations in the present study was 0.052, suggesting that a similar level of genetic differentiation occurred between the two populations in Ariake Bay, Japan.

For these two populations, the genetic differentiation within each population was maintained at a high level, and its variation was greater within the population consisting of individuals collected from the transplanted area (Fig. 4). There are three possible reasons why the variation in the degree of genetic differentiation within populations differed between the two populations. The first is that the degree of variation in genetic differentiation in these populations originally may have been different. Second, the methods of seedling production may lead to greater variation of genetic differentiation within populations in the transplanted area. In seedling production, broodstocks were randomly selected with closely related individuals collected from the same transplanted area or distantly related individuals collected from different transplanted areas. After stimulating the mature individuals to spawn, 20 to 30 individuals were placed in the same tank to produce fertilized eggs. Thus, the degree of genetic differentiation may have been more varied in the transplanted populations than in the wild habitat population. Third, the sample size may have been too small to accurately estimate the population. However, in the current situation in which the number of wild populations is small, it is difficult to verify the validity of a larger sample size.

### Guidelines concerning the best seed production and seed release methods

The current abundance of *S. constricta* is estimated to be very low based on the CPUE results of this study (Table 1). These results suggest that artificial seed production and release of seedlings could be an effective strategy for the recovery of *S. constricta* abundance in Ariake Bay, Japan. The present study suggested genetic differentiation between the two populations of *S. constricta* in Ariake Bay, and a high level of genetic differentiation within each population was maintained. Therefore, maintaining the two populations without interbreeding seems to be an appropriate conservation strategy. As of 2019, the Kashima lineage population is widely distributed in wild habitats, whereas the Nanaura lineage population is limited to the Takori mudflats. In order to avoid hybridization with the Kashima lineage population, the release of seedlings of the Nanaura lineage population should continue to be restricted to the mudflats within the current cove, with a transplanting area.

To produce seedlings to slow down the decline in genetic diversity within a population, mathematical models have evaluated five strategies: I) avoidance of line breeding, II) selective use of wild-born individuals, III) line breeding, IV) avoidance of artificial breeding of males, and V) selective use of wild-born males [29]. Among them, strategy II has been considered the most effective [29, 30]. The best way to produce seedlings of the Kashima lineage population would be to follow strategy II; all parental individuals would be randomly collected from wild habitats and previously produced seedlings would not be used as parents. In contrast, the Nanaura lineage population has already been maintained by a similar approach to strategy I; all parental individuals will be randomly collected from transplanted areas without excluding released seedlings. This approach gradually reduces the genetic diversity within the population. Indeed, lower pairwise *F*_*st*_ values were detected in the transplanted population (Fig. 4). Therefore, in the Nanaura lineage population, countermeasures should be taken to increase the number of parents used for seed production or to avoid using the produced seedlings as parents of the next generation.

In conclusion, we detected two genetically distinct populations of *S. constricta* in the Ariake Bay, Japan. Additionally, our results suggests that genetic differentiation within each population is maintained at a high level. In the future, monitoring the interbreeding status of the two genetically distinct populations and the genetic differentiation within each population is important to enhance the stock of *S. constricta* in Japan by producing and releasing seedlings, while preserving genetic diversity. As there was no significant difference in the morphology of the shells of these genetically distinct populations (Supplementary Fig. S1), DNA markers may be effective for discriminating between the two populations.

## Methods

### Sampling and genomic DNA extraction

Sampling of the razor clam *S. constricta* was conducted at 11 mudflats located along Ariake Bay (Fig. 1). Information on the sampling day and habitat types at each site is shown in Table 1. Habitats are divided into two types: transplanted area of artificial seedlings, and wild habitat. As mentioned above, artificial seedlings of *S. constricta* have been released into the transplanting areas established in natural seawaters since 1988. Therefore, artificially produced individuals live within the transplanted area, and the wild habitat would be populated by natural individuals and offspring of seedlings from the transplanted areas. The ages of individuals in the transplanted areas were estimated by calculating backwards from the year of release. At each site, razor clams were explored by hand and their abundance was assessed using the catch per unit effort index (CPUE), calculated as the number of clam individuals collected divided by the number of explorers and sampling time (i.e., ind. person^−1^ hour^−1^). The samples were stored at -80°C until genomic DNA extraction. In addition to the collected individuals, the oldest seedlings (produced in 2005, *n* = 2) stored at the Saga Prefectural Ariake Fisheries Research and Development Center, were included in the RAD-Seq analysis for comparison with seedlings produced in the past. Genomic DNA was extracted from finely chopped foot tissue of the specimens using DNAs-ici!-F (RIZO, Tsukuba, Japan), and the concentration of the DNA was assessed using a Qubit 3.0 Fluorometer (Thermo Fisher Scientific).

### RAD-Seq analysis

The preparation for *the Eco*RI RAD-Seq library has been described previously [28], based on the established method [31]. The library was sequenced with 150 bp paired-end reads in one lane of an Illumina HiSeq X (Illumina, San Diego, CA, USA). The RAD-Seq data were analyzed using the Stacks program (version 2.52) [32]. The data were quality filtered using the process_radtags program in the Stacks package (-r -c -q, -adapter_1, adapter_2, and adapter_mm2). Filtered sequences were registered in the DRA (DRA accession number: DRA010922). The data were aligned with the reference genome of *S. constricta* [33] (NCBI/DDBJ BioProject number: PRJNA508451) using BWA-MEM (version 0.7.17) [34]. Aligned data were analyzed using the ref_map.pl script of the Stacks package.

### Population genetics analysis

Principal component analysis (PCA) was conducted based on a variant call format (VCF) file using the SNPrelate program (v.1.18.1) [35]. The VCF file was created by the populations program of the Stacks package (--rm-pcr-duplicates, -R 0.7, --write-single-snp, --min-maf 0.05) [36]. The admixture analysis was conducted based on the PLINK file using ADMIXTURE (v. 1.3.0) [37]. The PLINK file was created with the same Stacks populations program that created the VCF file [38]. The genetic diversity of razor clams among and within populations was assessed using AMOVA *F*_*st*_ values obtained from ‘populations.fst_files’, which were calculated by the Stacks populations program. To visualize the distribution of all data points, pairwise *F*_*st*_ values between individuals in each population are shown in boxplot with beeswarm using R (package: ‘beeswarm’) [39].

### Phylogenetic tree analysis

Phylogenetic tree analysis was performed using two different approaches: the maximum likelihood (ML) method and SVDquartets analysis. Multiple alignments within the cluster were obtained using the Stacks package populations program mentioned earlier. The phylogenetic tree based on the ML method was constructed using the RAxML program (version 8.2.12) [40] (-f = a, -x = 12,345, -p = 12,345, -N (bootstrap value) = 1,000, and -m = GTRGAMMA). The SVDquartets analysis based on the coalescent model was computed as implemented in PAUP ver. 4.0a 167 using multiple alignments [41, 42]. The number of bootstrap analyses replicates was 1,000. In each analysis, the oldest samples (i.e., sample ID, 2005.1; 2005.2), were used as the root.

## Acknowledgments

The authors are grateful to the Saga Prefectural Ariake Fisheries Research and Development Center for supporting the field survey and providing samples and information on seedling production. This work was supported by the “Projects for sophistication of production and utilization technology supporting local agriculture and marine industry” from Saga University.

## Author contributions

R.O. designed and wrote the original manuscript draft. R.O. and Y.N. performed the RAD-Seq analysis and bioinformatic analysis. All authors reviewed drafts of the manuscript and approved its final version.

## Supplementary material

**Fig. S1.**
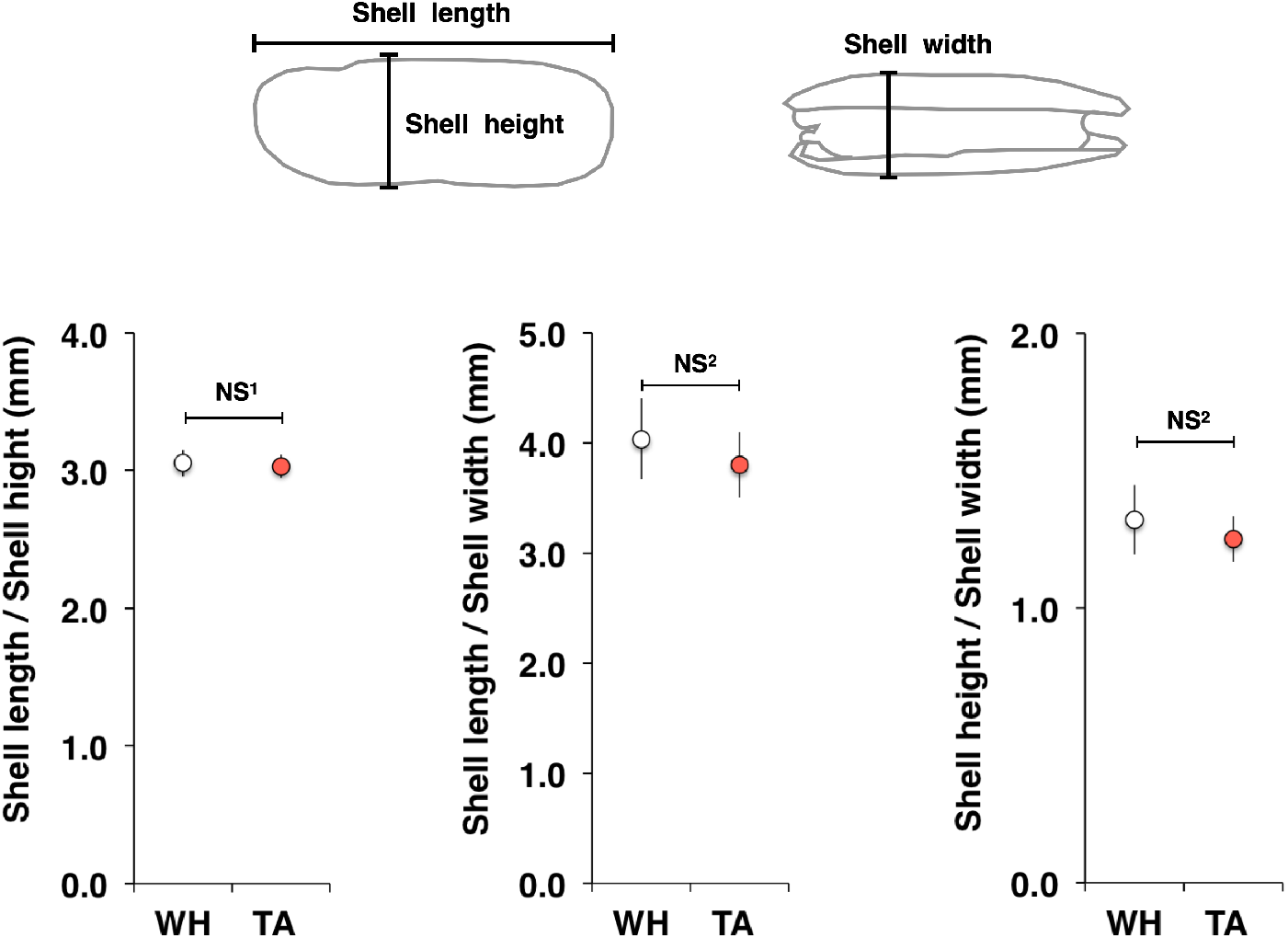
Comparison of shell morphology. WH: individuals in wild habitat (n=17), TA: individuals in transplanted area (n=11). NS^1^: non-significant difference between the comparison groups using the Student’s t test. NS^2^: non-significant difference between the comparison groups using the Wilcoxon rank sum test. Bar indicate the SD.

